# Modeling simian immunodeficiency virus (SIV) latency in primary rhesus macaque CD4^+^ T cells

**DOI:** 10.1101/2025.04.22.650029

**Authors:** Lea M. Matschke, Matthew R. Reynolds

## Abstract

Simian immunodeficiency virus (SIV)-infected rhesus macaques are valuable models for HIV cure research, offering insights into tissue reservoirs and testing reservoir-reduction strategies. Despite this utility, low frequencies of latently infected cells *in vivo* limit mechanistic studies of viral latency *ex vivo. In vitro* latency models have addressed this limitation for HIV, advancing our understanding of viral persistence. However, no comparable models exist for SIV. To address this gap, we developed an *in vitro* model of SIV latency in primary rhesus macaque CD4^+^ T cells, optimizing conditions to promote viral entry while maintaining cells in a minimally activated, non-proliferating state. After 12 days in culture, ∼1-3.3% of cells harbored SIV DNA, primarily as intact proviruses within central and transitional memory CD4^+^ T cell subsets. These cells remained quiescent, exhibiting minimal spontaneous viral protein production, but could be reactivated by potent T-cell stimulation and benchmark latency-reversing agents. Collectively, this model generates SIV-latently infected cells that resemble predominant cellular reservoirs *in vivo—*quiescent memory CD4^+^ T cells carrying inducible proviruses. This system provides a platform for investigating mechanisms of SIV latency, identifying shared and virus-specific features of HIV and SIV persistence, and evaluating strategies to reactivate or silence viral reservoirs.

**Importance:** Viral latency–HIV’s ability to persist in a dormant state within CD4^+^ T cells–remains a critical barrier to developing a cure. Rhesus macaques infected with simian immunodeficiency virus (SIV) are valuable models for studying HIV, but mechanistic studies of viral persistence are limited by the low frequency of latently infected cells *in vivo*. While *in vitro* models have advanced our understanding of HIV latency, comparable tools for studying SIV were lacking. To address this gap, we established a novel model of SIV latency using rhesus macaque CD4^+^ T cells, mirroring key features of natural reservoirs. This model provides a platform for studying how viral latency is established and maintained, conducting direct comparisons between HIV and SIV infections, and evaluating potential cure strategies.

## Introduction

*In vitro* models of human immunodeficiency virus (HIV) latency have been instrumental in identifying viral and cellular factors regulating latency (1). These models complement *ex vivo* studies using patient-derived latently infected cells, which are often too rare *in vivo* for detailed characterization (2−4). Similarly, simian immunodeficiency virus (SIV)-infected rhesus macaques are widely used to model HIV persistence, enabling longitudinal analysis of tissue reservoirs and preclinical testing of therapeutic interventions. Although HIV and SIV share many biological features, each has evolved host-specific adaptations that may shape how latency is established and maintained (5, 6). Despite the value of SIV/rhesus macaque models, primary cell-based *in vitro* latency models—such as those used for investigating HIV—have not yet been developed for SIV. Establishing such models would enable direct comparisons of the mechanisms governing HIV and SIV latency, strengthening SIV models as tools for HIV cure research.

*In vitro* latency models aim to replicate key features of HIV reservoirs, including the activation state and metabolic profile of naturally infected cells (1, 7, 8). In both antiretroviral therapy (ART)-treated humans and rhesus macaques, HIV/SIV reservoirs primarily consist of long-lived resting memory CD4^+^ T cells (9-13). Early models of HIV latency relied on immortalized cell lines with integrated proviruses, which provided crucial insights into the regulation of viral gene expression (14, 15). However, the high metabolic activity and continuous proliferation of these cells do not accurately reflect the quiescent nature of resting CD4^+^ T cells *in vivo*, limiting their relevance for investigating cell-intrinsic mechanisms of viral latency (1, 16).

To better reflect naturally infected cells, several groups developed HIV latency models using primary CD4^+^ T cells (7, 17-24). One such approach directly infects resting T cells (1). However, while more biologically relevant than immortalized cell lines, these direct infection models share a major challenge: quiescent CD4^+^ T cells are inherently resistant to retroviral infection and restrict key steps in virus replication, including viral entry, reverse transcription, and nuclear import (25, 26). To overcome these barriers, direct infection models typically treat resting T cells with cytokines (e.g., IL-2, IL-7, IL-15) (27-29) or chemokines (e.g., CCL19, CCL21) (30, 31) to increase their permissiveness, followed by infection via spinoculation to enhance virus-cell interactions (32, 33). While these models generate relatively low frequencies of latently infected cells (1, 7), their responses to mitogens and latency reversal agents (LRAs) resemble those of patient-derived CD4^+^ T cells (7, 31). However, despite their utility in advancing HIV research, the absence of comparable primary cell-based SIV latency models has limited detailed analysis of SIV latency and comparative studies across species and viral systems (1).

To address this gap, we developed an *in vitro* model of SIV latency using primary rhesus macaque CD4^+^ T cells. This approach primarily established latency in resting memory CD4^+^ T cells, which predominantly harbor HIV and SIV *in vivo* (12, 34). The cultured cells remained quiescent and harbored mostly intact proviruses that were inducible upon stimulation. This model bridges existing *in vitro* HIV latency models with *in vivo* SIV studies, providing a physiologically relevant tool to investigate mechanisms of SIV latency, evaluate responses to candidate LRAs and latency-promoting agents (LPAs), and directly compare HIV and SIV persistence.

## Results

### Modeling SIV latency in primary rhesus macaque CD4^+^ T cells

Our goal was to develop an *in vitro* method for generating latently infected cells using replication-competent SIV and primary rhesus macaque CD4^+^ T cells, building on direct infection models of HIV latency (27, 30). Specifically, we aimed to: (1) increase the susceptibility of resting CD4^+^ T cells to SIV infection, while (2) limiting cellular activation and proliferation, and (3) minimizing spontaneous SIV production.

Our first objective was to prime resting CD4^+^ T cells for SIV infection without inducing overt activation or proliferation. To achieve this, we sought to decouple viral permissivity from cellular activation by targeting the restriction factor sterile alpha motif and histidine-aspartate domain-containing protein 1 (SAMHD1). In quiescent T cells, SAMHD1 restricts infection by depleting cytoplasmic deoxynucleoside triphosphates (dNTPs), limiting their availability for viral DNA synthesis (25, 35). However, during the transition into the G_1_ phase of the cell cycle, cyclin-dependent kinases CDK1 and CDK2 phosphorylate SAMHD1, suppressing its antiviral activity (36, 37). We aimed to exploit this regulatory network by reversibly transitioning CD4^+^ T cells into the G_1_ phase through two complementary approaches: stimulating with IL-15 (27, 38) and inhibiting forkhead box protein O1 (FOXO1), a global transcription factor and positive regulator of cellular quiescence (39). Importantly, without additional proliferative signals, CD4^+^ T cells that enter G_1_ can revert to cellular quiescence (ie., G_0_ phase) once FOXO1 activity resumes (7, 22).

To implement this approach, we cultured resting CD4^+^ T cells with IL-15 and the small-molecule FOXO1 inhibitor AS1842856, along with a low concentration of IL-2 to support cell survival (Fig. 1). After five days, we infected the cells with clonal SIV via “magnetofection,” which uses magnetic nanoparticles to increase virus-cell interactions and improve infection efficiency (32). After infection, cells were cultured for two days without supplemental cytokines or antiretroviral drugs to foster proviral integration. At 2 days post-infection (dpi), we switched the cultures to an immunosuppressive medium containing the HIV/SIV protease inhibitor saquinavir to limit ongoing virus replication, along with cytokines TGF-β, IL-8, and IL-10 to suppress T cell activation and promote viral latency (40). After five additional days, we selectively depleted cells expressing the activation markers CD25 or HLA-DR, yielding the final population of cells.

**Fig 1.**
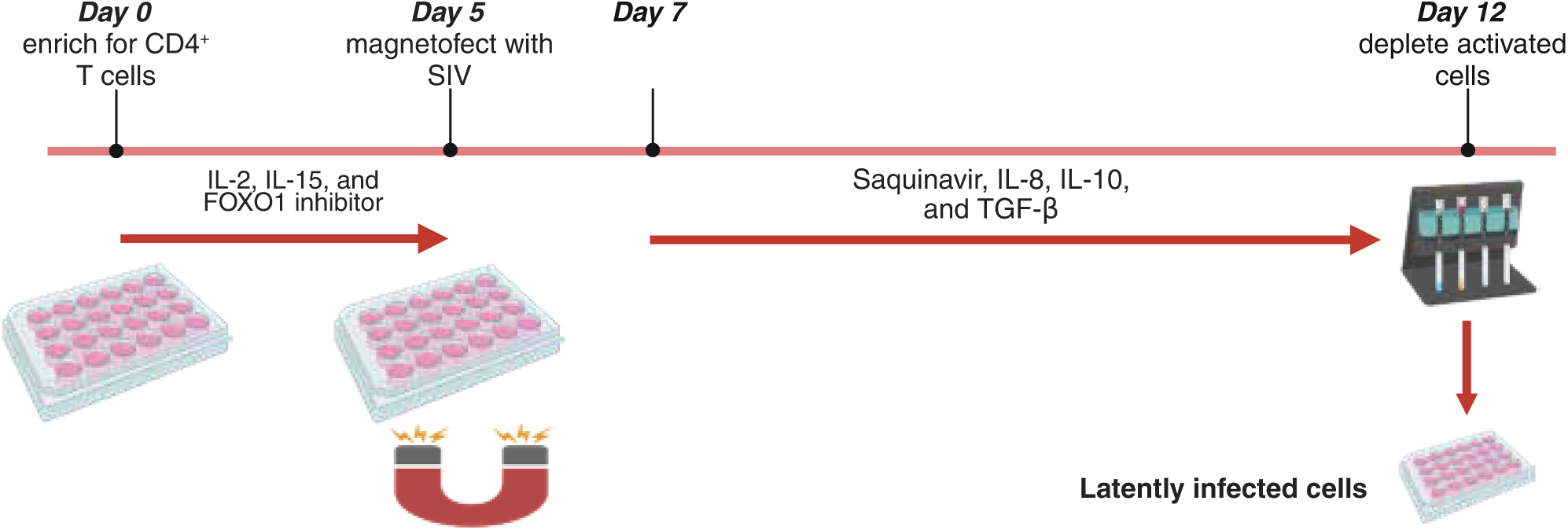
Generation of SIV latently infected cells in primary CD4^+^ T cells. CD4-expressing cells were enriched from PBMC and cultured from day 0 to 5 in R10 media supplemented with IL-15, IL-2, and a FOXO1 inhibitor (AS1842856). On day 5, cells were infected with SIV via magnetofection and cultured for 48 hours in standard R10 media. On day 7, the medium was replaced with R10 containing the protease inhibitor saquinavir and an immunosuppressive cytokine cocktail (IL-8, IL-10, and TGF-β). On day 12, cells expressing activation markers HLA-DR or CD25 were depleted via magnetic bead separation to generate a homogeneous population of resting CD4^+^ T cells. Created with https://BioRender.com.

### CD4^+^ T cells are maintained in a resting state

One of our goals was to maintain the CD4^+^ T cells in a quiescent, non-proliferating state throughout the culture period. Therefore, to determine whether IL-15 signaling and FOXO1 inhibition induced proliferation, we examined cell division at three key time points: immediately after CD4^+^ T cell isolation (day 0), at the time of infection (day 5), and at the end of the culture period following CD25^+^ and HLA-DR^+^ depletion (day 12). At each time point, we assessed DNA ploidy using the intercalating dye 7-AAD and monitored cell division based on the incorporation of bromodeoxyuridine (BrdU), a thymidine analog, over 20 hours. Cells were categorized as apoptotic (BrdU^-^/7-AAD^low^), dividing (BrdU^+^/7-AAD^intermediate/high^; S + G_2_ + M phases), or non-dividing (BrdU^-^/7-AAD^intermediate^; G_0_ + G_1_ phases) (Fig. S1A). At all three time points, ≥98.9% of live CD4^+^ T cells remained non-dividing, indicated by the absence of BrdU incorporation (mean±SD: 99.5±0.4) (Fig. 2A; Fig. S1B). In contrast, activated CD4^+^ T cell controls showed markedly higher BrdU incorporation and 7-AAD staining (Fig. S1A). To distinguish non-cycling T cells in G_0_ and G_1_, we measured intracellular Ki-67 expression, a marker of cellular proliferation that is expressed after G_1_ entry, with levels increasing through the S/G_2_/M phases (Fig. S1C). Consistent with minimal cell cycle progression, Ki-67 expression remained low and stable throughout the culture period (mean±SD: 0.9±0.1), with only 0.8±0.5% of live CD4^+^ T cells expressing Ki-67 on day 12 (Fig. 2B; Fig. S1D). Together, these results indicate that IL-15 signaling and FOXO1 inhibition did not induce T cell proliferation, as the cells maintained resting phenotypes throughout the culture period, consistent with previous studies (41-44).

**Fig 2.**
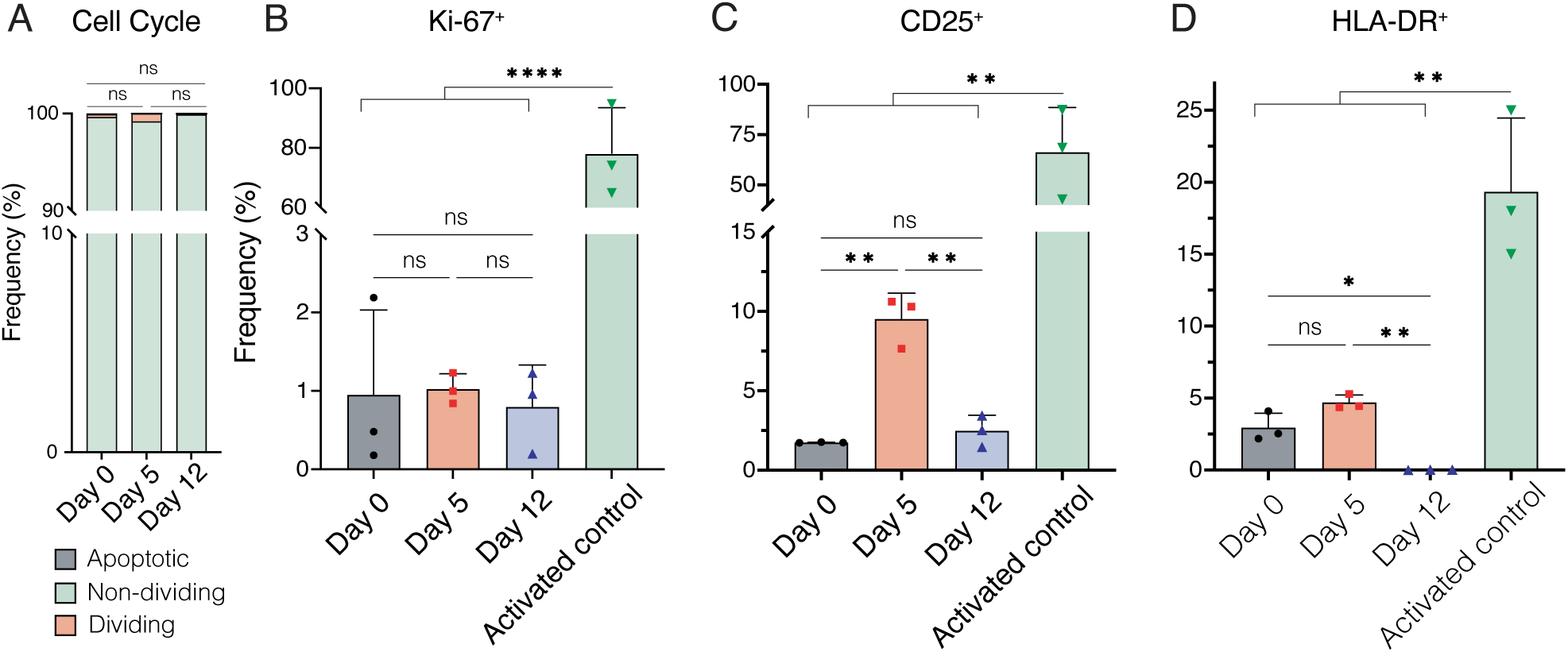
Cultured cells exhibit quiescent, non-activated phenotype. Cell cycle status and activation marker expression were assessed by flow cytometry at three timepoints: after CD4^+^ T cell isolation (day 0), at infection (day 5), and at the end of culture (day 12). (A) Cell cycle progression was determined by 7-AAD staining and BrdU incorporation over 20 hours to identify apoptotic (BrdU^-^/7-AAD^low^), non-dividing (G0 + G1 phase; BrdU^-^/7-AAD^intermediate^), or dividing (S + G2 + M phases; BrdU^+^/7-AAD^intermediate/high^) cells. (B-D) Expression of activation markers Ki-67 (B), CD25 (C), and HLA-DR (D) at each time point. Individual data points are shown as symbols, with boxes indicating the mean and error bars representing standard deviation. Data represent results from three independent replicates from the same donor macaque. For all panels, statistical analysis was performed using repeated-measures ANOVA: two-way for panel A and one-way for panels B-D, followed by Tukey’s post-hoc test. *p < 0.05; **p < 0.01; **** p < 0.0001.

We next evaluated whether the culturing conditions induced T cell activation by measuring CD25 and HLA-DR expression at the same three time points. By day 5, CD25^+^ frequencies increased significantly from day 0 (p=.004), though levels remained significantly lower than in activated controls (p=.003; Fig.2C). This modest ∼7.8% increase in CD25 expression from baseline aligns with previous reports of IL-15-mediated CD25 upregulation in human CD4^+^ T cells (27) and is within ranges previously reported for resting cells (28, 45-47). HLA-DR expression showed a modest, non-significant increase over the same period (p=.08, Fig. 2D). As expected, exposure to immunosuppressive cytokines followed by selective depletion of CD25^+^ and HLA-DR^+^ cells reversed these trends, returning to baseline frequencies by day 12 (Fig. 2C-D). Thus, while early culture conditions transiently upregulated activation markers, the final CD4^+^ T cell populations exhibited activation levels comparable to freshly isolated CD4^+^ T cells, supporting a resting cell phenotype (47).

### Cells harbor SIV DNA primarily within T_CM/TM_ cell subsets

Having established that our cells remained quiescent, we next evaluated their rates of infection by quantifying cell-associated SIV DNA (CA-vDNA) using droplet digital PCR (ddPCR). We measured SIV *gag* and normalized our results to the housekeeping gene ribonuclease protein p30 (*rpp30*) to express infection levels per cellular genome equivalents. On average, cultures contained ∼10,000 SIV *gag* DNA copies per million cell equivalents (eq.) (Fig. 3A) (mean±SD: 10,316±5,924; min-max: 1,630-18,674; 95% CI: 4,837-15,794), consistent with direct infection models of HIV latency (30, 31, 45). Infection rates varied between replicates and donors, reflecting both experimental and biological variability, a characteristic shared with primary cell models of HIV latency (7).

**Fig 3.**
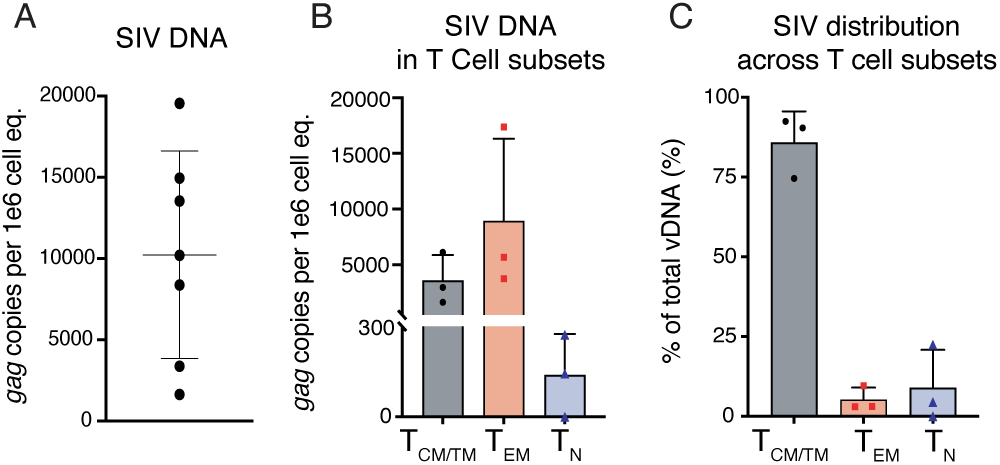
SIV preferentially infects memory CD4^+^ T cells. (A) Total cell-associated viral DNA (vDNA) in CD4^+^ T cells. Each point represents an independent infection replicate (n=7) from three rhesus macaques. (B and C) Live CD4^+^ T cells were sorted into central/transitional (T_CM_/T_TM_), naïve (T_N_), and effector memory (T_EM_) populations based on CD28 and CD95 expression. (B) vDNA content of each CD4^+^ T cell subset. (C) Relative contribution of each CD4^+^ T cell subset to the total vDNA pool based on both infection frequencies and relative abundance. Data represent the mean from three infection replicates from a single donor rhesus macaque. vDNA measurements were normalized to cell equivalents (eq) using the rhesus macaque *rpp30* gene. All ddPCR reactions were performed in duplicate with ≥3 technical replicates; error bars indicate standard deviations (SD).

We next determined the distribution of latently infected cells across major CD4^+^ T subsets by sorting live CD4^+^ cells into three populations: T_N_ (CD28^+^ CD95^-^), T_CM/TM_ (CD28^+^ CD95^+^), and T_EM_ (CD28^-^ CD95^+^) (Fig. S2B). After sorting, we quantified CA-vDNA in each subset. Consistent with *in vitro* models of HIV latency, SIV DNA was most abundant on a per-cell basis in T_EM_ cells (mean±SD: 9,451±7,084 *gag* copies per million cell eq.; min-max: 3,759-17,384) (Fig. 3B), potentially reflecting their heightened metabolic activity (48). However, since T_EM_ cells constitute only a small fraction of peripheral CD4^+^ T cells (49) (Fig. S2B), they accounted for only ∼6% of total infected populations (Fig. 3C). In contrast, T_N_ cells, which are abundant in the blood, harbored low levels of SIV DNA (mean±SD:139±136 copies per million cell eq.; min-max: 0-273) (Fig. 3B), but, due to their relative abundance in the cultures, represented ∼9% of total SIV-infected cells (Fig. 3C). As expected, the majority of latently infected cells resided within the T_CM/TM_ subsets (Fig. 3B-C) (mean±SD: 3,576±2,305 copies per million cell eq.; min-max: 1,635-6,124), representing ∼86% of all SIV infected CD4^+^ T cells, consistent with their contribution to latent SIV reservoirs in ART-suppressed rhesus macaques (50).

### SIV proviruses are primarily intact

Quantifying SIV *gag* alone provides valuable insights into infection rates but may overestimate the frequency of replication-competent infections by including unintegrated or defective proviruses, such as incomplete reverse transcripts or hypermutated genomes. To address this limitation, we employed the intact proviral DNA assay (IPDA), which selectively quantifies intact proviruses, excludes those with common defects, and corrects for sequence shearing during DNA extraction (51, 52). This assay offers a more comprehensive assessment of the proviral DNA landscape than measuring SIV *gag* alone, allowing us to distinguish between intact—potentially replication-competent proviruses—and defective viral genomes.

Our analysis revealed that cultured CD4^+^ T cells contained an average of ∼16,000 intact proviral copies per million cell eq. (mean±SD: 16,420±7,211; min-max: 6,908-24,215; 95% CI: 4,945-27,894), accounting for ∼66% of total SIV DNA (Fig. 4) and suggesting that high proportions of proviral genomes were replication-competent. In contrast, defective proviral DNA—characterized by 5’ or 3’ deletions and/or hypermutations— comprised only ∼24% of total SIV DNA. Specifically, 5’ deletions averaged 2,207±1,191 copies per million cell eq. (min-max: 457-3,111; 95% CI: 312-4,103) and 3’ deletions averaged 3,827±1,156 copies per million cell eq. (min-max: 2,462-5,289; 95% CI: 1,988-5,666). Finally, unintegrated 2-long terminal repeat (2-LTR) circles made up ∼9% of the total SIV DNA (mean±SD: 2,247±1,219 copies per million cell eq.; min-max: 622-3,552; 95% CI: 307-4,187). These results demonstrate that our model efficiently establishes latency with predominantly intact and potentially inducible proviruses.

**Fig 4.**
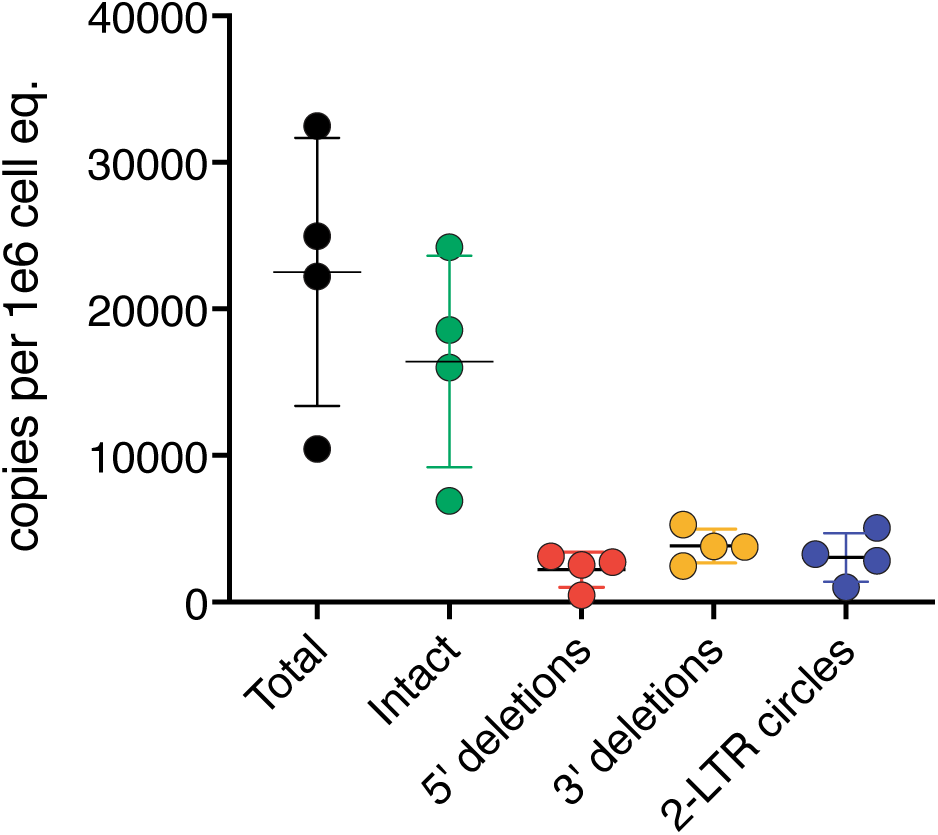
Intact SIV proviral DNA predominates in latently infected cells. IPDAs were performed to quantify different forms of viral DNA in latently infected cells. Results show copies per million cell equivalents (eq.) of total viral DNA (vDNA), intact vDNA, proviruses with 5’ and 3’ deletions, and 2-LTR circles across four independent cultures from the same donor macaque. Individual data points are shown as dots, with bars indicating the mean and error bars representing standard deviation. vDNA copy numbers were normalized to cell equivalents using the rhesus macaque *rpp30* reference gene. All ddPCR reactions were performed in duplicate with a minimum of three technical replicates.

### SIV latently infected cells contain reactivatable virus

Building on these findings, we next investigated whether these proviruses were inducible. While the SIV IPDA distinguishes between intact and defective proviruses, additional factors—such as proviral integration sites, epigenetic modifications, and host transcription factor availability—may influence viral reactivation (53). To determine whether the proviruses in our model were functional, we assessed their ability to produce viral proteins upon potent stimulation. We incubated the cells with anti-CD2/CD3/CD28-coated T cell activation beads for 36 hours or left them untreated as controls, then measured intracellular SIV Gag. Unstimulated cells showed minimal spontaneous virus production, with SIV Gag expression consistently below 0.12% of live T cells (Fig. 5A). In contrast, upon stimulation 0.2–0.6% of live T cells expressed SIV Gag (Fig. 5B), a level comparable to HIV direct infection models (30, 31). This increase in viral protein expression indicates that our system generates SIV-infected cells that rarely express viral proteins when unstimulated but readily reactivate in response to strong stimuli.

**Fig 5.**
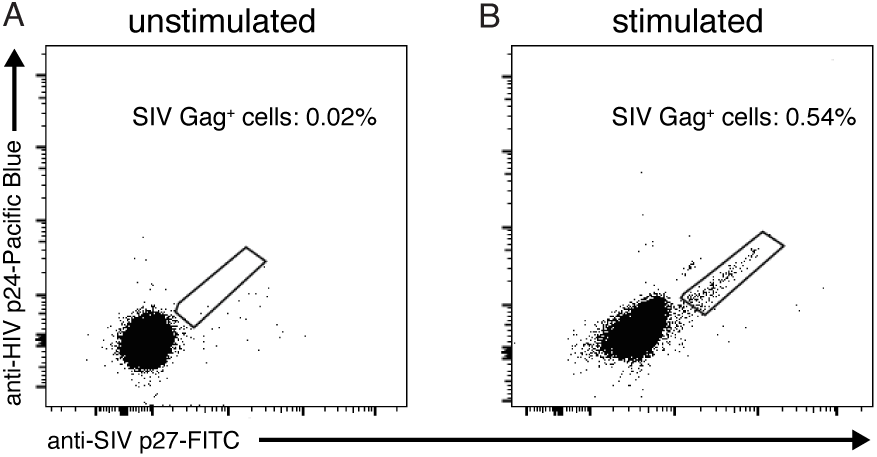
T cell activation triggers SIV reactivation from latency. We collected cells on day 10.5 of the cultures after depleting cells expressing activation markers CD25 or HLA-DR. Cells were either (A) left unstimulated or (B) stimulated with anti-CD2/CD3/CD28-coated beads for 36 hours. Flow cytometry dot plots show intracellular SIV Gag detection using two antibodies binding distinct epitopes: anti-SIV p27-FITC (clone 55-2F12, x-axis) and anti-HIV p24-Pacific Blue (clone 183-H12-5C, y-axis). These representative analyses used cells from a single *in vitro* latency culture, with gating on singlets/lymphocytes/live/CD3^+^.

We next evaluated how our latently infected cells responded to benchmark LRAs with distinct mechanisms of action: IL-15 (PI3K/Akt activator), SAHA (histone deacetylase inhibitor), bryostatin-1 (protein kinase C agonist), tumor necrosis factor α (TNF-α; NF-κB activator), and ionomycin (calcium ionophore; NFAT activator). Cultured cells were exposed to individual LRAs for 48 hours in the presence of raltegravir, an integrase inhibitor that prevents ongoing viral replication. SIV Gag levels in culture supernatants were then quantified by ELISA across three independent experiments and normalized to unstimulated (baseline) and bead-stimulated (maximum induction) controls.

Our results revealed differential responses to these LRAs (Fig. 6). IL-15, bryostatin-1, and ionomycin significantly increased virion production, whereas TNF-α and SAHA had minimal effects. We observed that the effectiveness of these agents corresponded to their mechanisms of action. The most potent LRAs activate transcription factors that facilitate the recruitment or activation of positive transcription elongation factor b (p-TEFb) at the viral promoter. Specifically, bryostatin-1 activates NF-κB, ionomycin modulates NFAT, and IL-15—the strongest inducer—can affect both pathways, leading to p-TEFb-mediated transcription of viral genes (27, 54, 55). Despite its known role in NF-κB activation, TNF-α was ineffective in activating latently infected cells generated by our model, possibly due to low TNF-α receptor expression on peripheral memory CD4^+^ T cells, as previously reported in primary cell cultures and HIV-infected individuals (7, 24). These response patterns align with observations from HIV direct infection models and highlight the central role of NF-κB and NFAT signaling in SIV reactivation (7, 56, 57). Collectively, these findings demonstrate that our *in vitro* model generates SIV latently infected cells that respond differentially to LRAs in a physiologically relevant manner, making it a valuable tool for investigating mechanisms of SIV latency and evaluating candidate latency reversal strategies.

**Fig 6.**
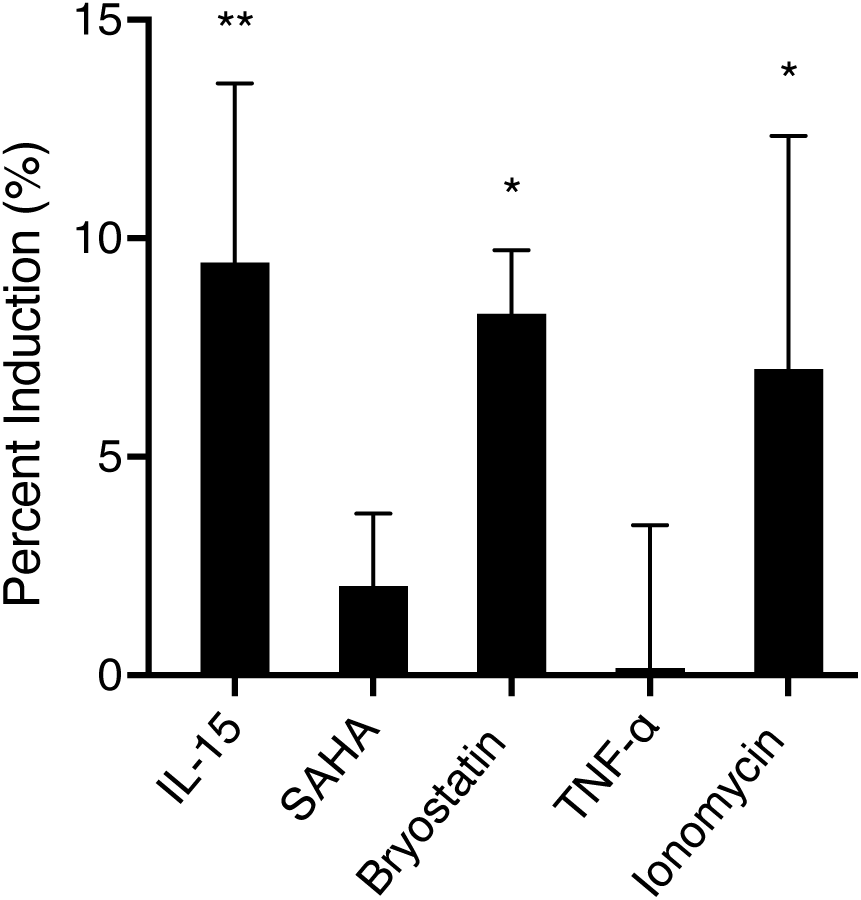
Latency reversal agents induce SIV reactivation. We collected cells on day 10.5 of the cultures after depleting activated cells and treated them with benchmark LRAs plus the integrase inhibitor raltegravir. After 48 hours, we measured extracellular SIV p27 in culture supernatants by ELISA. Data are normalized across three independent infection replicates from a single donor, with unstimulated controls set as 0% and anti-CD2/CD3/CD28 bead stimulation defined as 100% activation. Statistical significance was calculated by 1-way ANOVA with Dunnett’s multiple comparisons test comparing each LRA to unstimulated controls. Error bars represent standard deviation. *, p<0.05; **, p<0.01.

## Discussion

In this study, we describe an *in vitro* model of SIV latency using primary rhesus macaque CD4^+^ T cells, bridging existing HIV latency models and *in vivo* macaque studies. Our approach reliably establishes latent infections in resting memory CD4^+^ T cells, recapitulating key features of *in vivo* HIV/SIV reservoirs. These cells predominantly harbor intact proviruses that remain inducible upon stimulation. Altogether, this model provides a valuable platform to investigate mechanisms of viral latency and evaluate HIV cure strategies in a physiologically relevant system that bridges the gap between *in vitro* human models and *in vivo* macaque studies.

A major challenge in developing this model was overcoming the intrinsic resistance of quiescent CD4^+^ T cells to SIV infection. Although both HIV and SIV can infect resting T cells (38, 58), cellular quiescence presents multiple barriers to infection, including a rigid actin cytoskeleton and low levels of cytosolic dNTPs and ATP, which collectively hinder viral entry, reverse transcription, and nuclear import (25, 56, 59-62). Among these restrictions, SAMHD1 blocks reverse transcription by continuously depleting cytosolic dNTPs (25). However, as resting cells enter the G_1_ phase, CDK1 and CDK2 phosphorylate and inhibit SAMHD1 activity, increasing dNTP availability for viral DNA synthesis (36).

Therefore, we reasoned that treating cells with IL-15 and the FOXO1 inhibitor AS1842856 would enhance SIV infection of resting cells, as both have been shown to enhance Cdk1 activity in primary human T cells (22, 27). IL-15 can further support infection through the mTOR pathway, boosting cellular metabolism and expanding dNTP pools needed for reverse transcription (63). Beyond supporting reverse transcription, these treatments may also promote viral entry and nuclear trafficking through increased ATP production and actin cytoskeleton remodeling (38, 43, 64, 65). Collectively, these mechanisms likely promoted our relatively efficient SIV infection of resting CD4^+^ T cells.

A key aspect of any latency model is its response to LRAs. These agents must overcome viral and cellular factors that govern retroviral latency, including proviral integration sites, chromatin accessibility, epigenetic modifications, transcription factor availability, and Tat expression (53). When we tested cells from our model with benchmark LRAs targeting different cellular pathways, we observed that compounds modulating host transcription factors effectively reactivated SIV, whereas the histone deacetylase (HDAC) inhibitor SAHA had minimal impact. These distinct LRA responsiveness patterns, consistent with previous studies (7, 66, 67), indicate that transcription factors such as NF-κB and NFAT play a more prominent role in reversing latency in our model than epigenetic remodeling alone. Notably, this LRA response profile closely resembled those observed in HIV direct infection models and patient-derived latently infected cells (7, 23), demonstrating our model’s relevance for comparative studies between HIV and SIV latency. Future studies in our system could explore novel LRAs, combinatorial LRA treatments to enhance viral reactivation, or LPAs to promote deeper latency (66-70).

Beyond screening therapeutic agents, our latency model could support studies determining how reservoir size affects the timing of viral rebound and the levels below which durable immune control becomes achievable. By infusing autologous latently-infected cells into SIV-naïve rhesus macaques, it may be possible to generate reservoirs containing known quantities of intact proviruses. This approach could provide insights into how physiological factors—including host immune responses—affect reservoir maintenance and viral reactivation, helping identify minimum reservoir sizes for sustained viral control. More broadly, this system could facilitate direct comparisons of the efficacy of therapeutic agents *in vitro* and *in vivo* using the same population of latently infected cells.

Despite these potential applications, several aspects of our approach could be refined to better reflect the full spectrum of *in vivo* viral reservoirs and improve overall model performance. One challenge is that our model generates relatively few latently infected effector memory T cells, which can be a sizable component of *in vivo* reservoirs (71). This underrepresentation is due to the low frequency of effector memory T cells among peripheral CD4^+^ T cells—our starting material—which are primarily composed of naïve and central memory T cells (72). Enriching for memory CD4^+^ T cells at the start of the culture period could help address this limitation; however, rhesus macaques lack naïve T cell markers equivalent to human or murine CD45RA, complicating the selective depletion of naïve cells (72). Additionally, while incubating resting CD4^+^ T cells with IL-15 and a FOXO1 inhibitor likely increased their permissivity to SIV infection, infection rates were still low relative to those observed in post-activation latency models (1, 8, 17, 19, 20). One potential approach to improve infection efficiency is to supplement the culture medium with exogenous deoxynucleosides (dNs) to increase intracellular dNTP concentrations, thereby aiding reverse transcription and integration (35, 73). Future refinements addressing these limitations could enhance the model’s utility for investigating mechanisms of viral latency and reactivation, strengthening its translational relevance.

In conclusion, we have developed a direct infection model of SIV latency using primary rhesus macaque CD4^+^ T cells, offering a physiologically relevant system for studying viral persistence *in vitro*. Our approach generates infected cells that closely resemble *in vivo* latent reservoirs, with latency predominantly established in resting CD4^+^ T_CM/TM_ cells that harbor intact, inducible proviruses. This model provides a valuable tool for direct comparisons between HIV and SIV latency, enabling identification of both shared and virus-specific mechanisms of persistence. It also provides a preclinical platform for evaluating novel therapeutic agents across species, enhancing the translational relevance of SIV/rhesus macaque models for HIV cure research.

## Materials & Methods

### Animal handling & care

Rhesus macaques (*Macaca mulatta*) were maintained at the Wisconsin National Primate Research Center (WNPRC) in accordance with guidelines approved by the University of Wisconsin-Madison Graduate School Institutional Animal Care & Use Committee (IACUC protocol G006049) and recommended in the National Research Council’s Guide for the Care and Use of Laboratory Animals’ Weatherall Report. Staff administered anesthesia and/or fluid replacement therapies as needed.

### Production of SIV stocks

We generated SIV_mac239_ stocks from hemi-genome plasmids (NIH AIDS Research and Reference Reagent Program), as described previously (74). Briefly, we seeded 5 x 10^5 Vero cells into individual wells of a 6-well tissue culture plate and cultured them overnight in R10 media (RPMI 1640 supplemented with 10% fetal bovine serum (FBS), 1% antibiotic-antimycotic solution, and 1% L-glutamine; all from Cytiva except the FBS, which was from Corning) at 37°C with 5% CO_2_. The following day, we transfected the Vero cells with SIV_mac239_-encoding plasmids using Lipofectamine (Invitrogen) and added CEMx174 cells to the cultures 24 hours later. After two additional days, we transferred the CEMx174 cells to T75 flasks and monitored them for syncytia formation. Upon observing widespread syncytia, we harvested cell-free supernatants daily for three consecutive days. The virus-containing supernatants were then divided into 1mL aliquots and stored in liquid nitrogen.

### *In vitro* latency model

#### Overview

We developed a 12-day protocol to generate latently SIV-infected CD4^+^ T cells isolated from rhesus macaque peripheral blood, consisting of cell isolation and priming, virus preparation, SIV infection via magnetofection, immunosuppressive conditioning, and depletion of activated cell populations.

#### Timeline

Days 0-5: Cell priming; Day 5: Infection; Day 7-12: Immunosuppressive conditioning; Day 12: Deplete cells expressing activation markers.

i. **CD4^+^ T cell isolation and priming:** We obtained PBMCs from whole blood by density-gradient centrifugation with Ficoll-paque PLUS (Cytiva). CD4^+^ T cells were isolated from PBMCs using a nonhuman primate CD4^+^ T cell isolation kit (Miltenyi Biotec) according to manufacturer’s instructions. CD4^+^ T cells were resuspended in Priming Media at 5 x 10^6-8 x 10^6 cells/mL and seeded in flat-bottom 48-well tissue culture plates (Corning) at 1 mL per well. Priming Media consisted of R10 supplemented with recombinant human IL-2 (10U/mL; R&D Systems), recombinant human IL-15 (10ng/mL; R&D Systems), and FOXO1 inhibitor AS1842856 (500nM; Sigma Aldrich). After 2-3 days, we replaced at least 500 µL in each well with fresh Priming Media. Cells were maintained at 37°C with 5% CO_2_ throughout the 12-day culture period.
ii. **Virus stock purification:** Before use, we purified SIV stocks using a 20% sucrose cushion to remove residual proteins (74). Briefly, virus stocks were layered onto 100 µL of 20% sucrose solution and centrifuged at >50,000xg for 1.5-3 hours at 4°C. After discarding supernatants, viral pellets (∼30-100µL) were diluted with ∼150 µL of phosphate**-**buffered saline (PBS) at room temperature.
iii. **SIV infection:** On day 5, we replaced Priming Media with equal volumes of R10 and infected cultured cells with SIV by magnetofection (32). First, 150-250 µL purified virus in PBS was added to 30 µL Viromag R/L transduction reagent (OZ Biosciences) at room temperature. After 15 minutes, this mixture was added to cultured cells and centrifuged at 670xg for 5 minutes. We then incubated the pelleted cells on Super Magnetic Plates (OZ Biosciences) for 1-3 hours at 37°C before resuspending.
iv. **Immunosuppressive conditioning:** At 2 days post-infection (day 7), we replaced R10 with equal volumes of Immunosuppressive Media, consisting of R10 supplemented with saquinavir mesylate (5µM; Sigma Aldrich), recombinant human TGF-β1 (10ng/mL; R&D Systems), recombinant human IL-8/CXCR8 (50ng/mL; R&D Systems), and recombinant human IL-10 (10ng/mL; Peprotech). Between days 9 and 10, we replenished ≥500 µL of Immunosuppressive Media in each well.
v. **Depletion of cells expressing activation markers:** To obtain final populations, we depleted activated (ie., CD25^+^ or HLA-DR^+^) subsets on day 12 of the culture period. Briefly, we incubated cells with biotinylated anti-CD25 (clone BC96; Biolegend) and anti-HLA-DR (clone LN3; Biolegend) monoclonal antibodies for 15 minutes at 4°C. Once labeled, cells were incubated with anti-biotin microbeads (Miltenyi Biotec) for 15 minutes at 4°C and passed through LS columns (Miltenyi Biotec) according to the manufacturer’s instructions. Final populations were collected from negative fractions.

### Flow cytometry

i. **General:** To detect cell surface markers, we stained cells with viability dye and fluorophore-conjugated antibodies at room temperature for 30-45 minutes. After two PBS washes, cells were fixed with 2% paraformaldehyde (PFA) in PBS for >10 minutes at room temperature. For intracellular staining, fixed cells were subsequently washed with FACS buffer (2% FBS in PBS) and permeabilized in Medium B Permeabilization Buffer (Invitrogen) with corresponding antibodies for 20 minutes at room temperature. Cells were then washed twice with FACS buffer and fixed with 2% PFA. For cell cycle analysis, we added 7-amino-actinomycin (7-AAD) intercalating dye after the final wash, according to manufacturer’s instructions. Data were collected on a Becton Dickinson (BD) FACSymphony A3 flow cytometer with FACSDiva acquisition software and analyzed with FlowJo v10 (TreeStar Inc.). Gating strategies are shown in Fig. S2.
ii. **Cell surface staining:** Cells were labeled with Near-IR fixable dead cell dye (Invitrogen) and antibodies against the following surface markers: CD3-Alexa Fluor 700 (clone SP34-2, BD), CD4-APC (clone OKT4, Biolegend), CD28-PE (clone CD28.2, BD), CD95-PE-Cy5 (clone DX2, BD), CCR7-Brilliant Violet 605 (clone G043H7, Biolegend), CD25-Brilliant Violet 711 (clone M-A251, Biolegend), and HLA-DR-PE/Dazzle594 (clone L243, Biolegend).
iii. **Cell cycle analysis:** We assessed BrdU uptake, DNA ploidy, and activation marker expression on days 0, 5, and 12 of the protocol. For comparison, primary CD4^+^ T cells were cultured in parallel with recombinant human IL-2 (70U/mL; R&D Systems) and nonhuman primate T cell activation beads (Miltenyi Biotec) at 0.6 beads/cell (Fig. S2A-C). Prior to each time point, we cultured 1x10^5–1x10^6 cells with and without BrdU for 20 hours. Cells were then stained with Near-IR fixable dead cell dye (Invitrogen) and antibodies against CD4-APC (clone OKT4, Biolegend), CD25-Brilliant Violet 421 (clone M-A251, Biolegend), HLA-DR-PE/Dazzle594 (clone L243, Biolegend), Ki-67-Brilliant Violet 421 (clone B56, BD), as well as BrdU-FITC and 7-AAD (both from the FITC BrdU Flow Kit, BD).
iv. **Intracellular SIV Gag staining:** On day 10.5, final populations were divided into stimulated or unstimulated groups and cultured at <8 x 10^6 cells/mL for the remainder of the culture period. For unstimulated groups, we replaced culture media with fresh Immunosuppressive Media without saquinavir to permit virus replication. Stimulated groups were cultured in R10 with raltegravir (50µM; SelleckChem) and T cell activation beads (Miltenyi Biotec) at 0.6 beads/cell. After 36 hours at 37°C with 5% CO_2_, cells were labeled with viability dye and antibodies against surface markers as well as anti-SIV Gag p27-FITC (clone 55-2F12) and anti-HIV Gag p24-Pacific Blue (clone 183-H12-5C), which recognize distinct epitopes on SIV Gag p27. The anti-HIV Gag p24 monoclonal antibody was isolated from hybridoma (ARP-1513; NHP AIDS Reagent Resource) supernatants and conjugated to Pacific Blue using a commercial kit (Invitrogen), following manufacturer’s instructions. The cells were gated on singlets/lymphocytes/live/CD3^+^/p24^+^/p27^+^.

### Cell sorting

To isolate distinct memory and naive T cell populations (72), we labeled the final cell preparations with Near-IR fixable dead cell stain (Invitrogen) and antibodies against CD4-APC (clone OKT4, Biolegend), CD28-PE (clone CD28.2, BD), and CD95-PE-Cy5 (clone DX2, BD) antibodies for 30 minutes at room temperature. After two PBS washes, we sorted live CD4^+^ cells into three subsets using a BD FACSJazz Cell Sorter: naïve (T_N_; CD28^+^/CD95^-^), central/transitional memory (T_CM/TM_; CD28^+^/CD95^+^), and effector memory (T_EM_; CD28^-^/CD95^+^) (Fig. S2 A-B).

### DNA extraction

We extracted genomic DNA from cultured CD4^+^ T cells one of two Qiagen kits: the DNeasy Blood & Tissue Kit for SIV *gag* quantification or the QIAmp DNA Mini Kit for the IPDA, as previously described (51, 75). Excess sample eluent was removed using the Vacufuge Plus System (Eppendorf), and DNA concentrations were measured using a Nanodrop One spectrophotometer (ThermoFisher).

### SIV *gag* ddPCR

Cell-associated viral DNA (CA-vDNA) was measured by SIV *gag* ddPCR. To prepare for the assay, we digested sample DNA with EcoRI-HF (New England Biolabs) per the manufacturer’s instructions. SIV *gag* and host *rpp30* were quantified in separate singleplex ddPCRs on the Bio-Rad QX200 ddPCR System, as previously described (75, 76) . Briefly, each 20 µL reaction contained 3.2 µL digested DNA (≤1,000ng), ddPCR Supermix for Probes (no dUTP) (Bio-Rad), 900nM primers, and 250nM probes (all from Integrated DNA Technologies (IDT)) (Table S1).

To prevent oversaturation of positive droplets during *rpp30* amplification, we diluted sample DNA for this target separately at fixed ratios to yield 0.5-2 ng/µL before transferring equal volumes to each 20 µL reaction. Dilution factors were then applied to *rpp30* frequencies to represent equivalent DNA input across all reactions.

For thermal cycling conditions, SIV *gag* reactions, droplets were cycled as follows: 95°C for 10min; 40 cycles of 94°C for 30s and 58°C for 60s; 98°C for 10min; 4°C hold. For singleplex *rpp30* ddPCRs: 95°C for 5min; 40 cycles of 94°C for 15s, 55°C for 30s, 72°C for 30s; 72°C for 5min; 4°C hold. Droplets were analyzed on the QX200 Droplet Reader with Quantasoft Studio Software (Bio-Rad). Data represent duplicate wells averaged across 2-3 replicates. Negative controls included no template controls and uninfected DNA from each macaque. As a positive control, we extracted DNA from 3D8 cells, an SIV-infected cell line (NHP AIDS Reagent Resource).

### IPDA

Intact and defective SIV DNA were quantified using the previously described intact proviral DNA assay (IPDA) as previously described (51, 77). Briefly, the SIV IPDA utilizes three duplexed ddPCR reactions targeting the following amplicons: (1) SIV *pol* and *env*(52), (2) SIV 2-LTR junctions (78) and *env*, and (3) two regions of the *rpp30* housekeeping gene (Table S2). For each reaction, corrected data were averaged from duplicate wells across ≥3 replicates. Intact proviruses were defined as *pol*^+^/*env*^+^ sequences without 2-LTR junctions, while total SIV DNA was calculated as the sum of *pol*^+^/*env*^+^, *pol*^+^/*env*^-^, and *pol*^-^/*env*^+^ sequences, irrespective of 2-LTR junctions (51, 79).

### SIV reactivation with LRAs

To assess the responsiveness of our latently infected cells to latency reversal agents (LRAs), we compared SIV Gag p27 production in culture supernatants after exposure to benchmark LRAs for 48 hours. Briefly, final populations were resuspended in R10 with raltegravir (50µM; SelleckChem, Houston, TX) and divided equally (1.2-2.5 x 10^6 cells/well) into individual wells of a 96-well plate at 225 µL/well. Parallel cultures were left unstimulated or exposed to individual LRAs for 48 hours, then SIV Gag p27 was quantified in culture supernatants using a commercial ELISA kit (Zeptometrix) according to the manufacturer’s instructions. Each replicate compared SIV p27 produced by unstimulated controls to the following stimuli: SAHA (250nM; MilliporeSigma), recombinant human IL-15 (25ng/mL; R&D Systems), bryostatin-1 (25ng/mL; R&D Systems), recombinant rhesus macaque TNF-α (10ng/mL; R&D Systems), ionomycin (750nM; Research Products International), and T cell activation beads (0.6 beads/cell).

To normalize across infection replicates, baseline (unstimulated) SIV p27 concentrations were subtracted from treatment values, and the differences were divided by maximum response (T cell activation beads) to yield percent reactivation. Data represent duplicate wells averaged across three infection replicates [n=3] from a single donor.

### Statistical Analysis

Data were analyzed using Prism 10.4 software (GraphPad) by one- or two-way repeated measures (RM) ANOVA with multiple comparisons. All data passed preliminary tests for single-pooled variances. For the analysis of Ki67, CD25, and HLA-DR expression, multiple comparisons were calculated with and without activated controls. Pairwise comparisons to activated controls were performed using the entire data set. For all other comparisons, activated controls were excluded from the analysis to increase statistical power for detecting differences between experimental groups.

## Acknowledgements

This study was supported by National Institutes of Health (NIH) grant R01AI174940. Additional support was provided by the Office of Research Infrastructure Programs/OD via grant P51OD011106 awarded to the WNPRC at the University of Wisconsin-Madison. The funders had no role in study design, collection, analysis, data interpretation, or manuscript preparation. The content is solely the responsibility of the authors and does not necessarily represent the official views of the NIH. The authors have no competing interest to declare.

The following reagent was obtained through the NIH HIV Reagent Program, Division of AIDS, NIAID, NIH: Anti-Human Immunodeficiency Virus 1 (HIV-1) p24 Hybridoma (183-H12-5C), ARP-1513, contributed by Dr. Bruce Chesebro and Dr. Hardy Chen. Additionally, Andrea Weiler and Dr. Thomas Friedrich (WNPRC Virology Services) assisted with the production of clonal virus stocks, and Kim Weisgrau (WNPRC Immunology Services) conducted FACS of memory and naïve populations.

