## Supplemental Figure 1 for "Modeling simian immunodeficiency virus (SIV) latency in primary rhesus macaque CD4^+^ T cells"

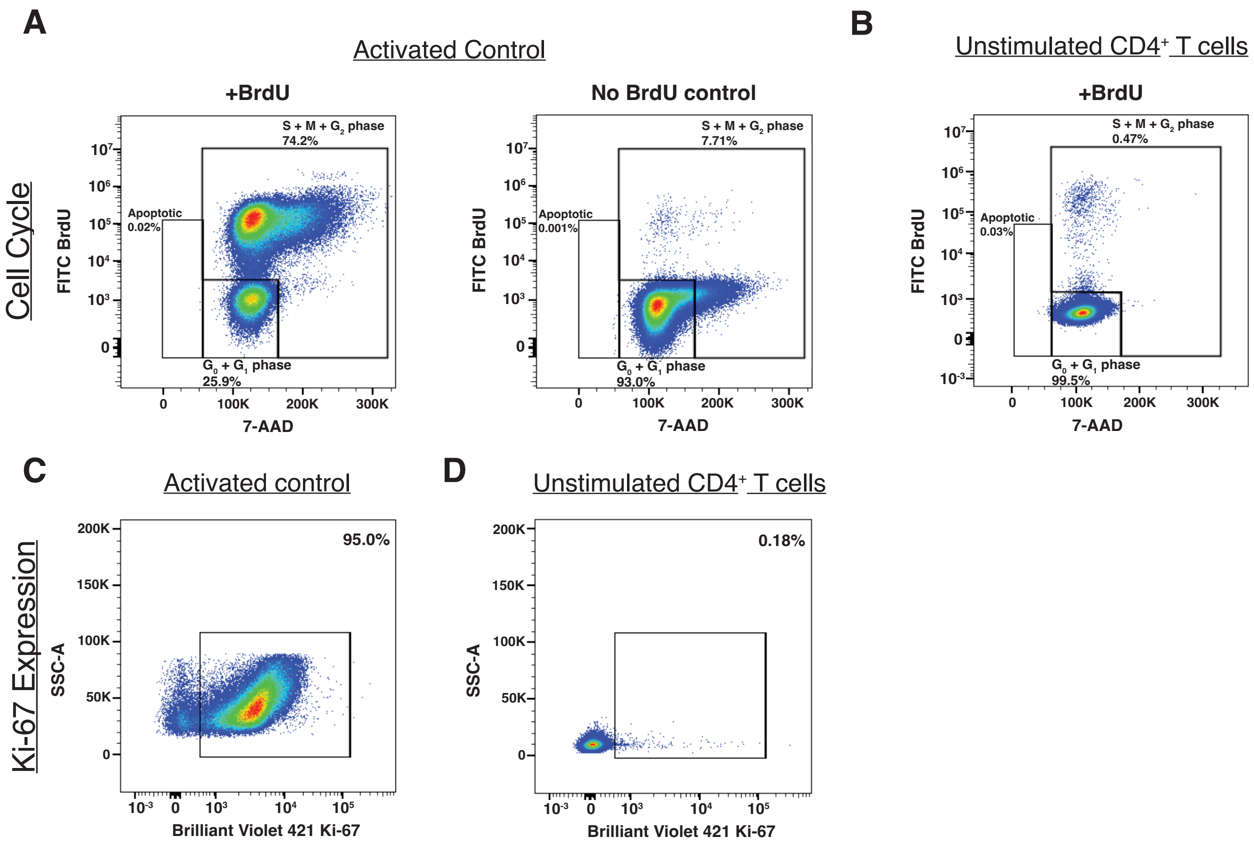


**Figure S1.** Representative flow cytometry analysis of cell cycle and proliferation marker expression. (A) Cell cycle analysis of CD4$^{+}$ cells incubated with T cell activation beads. The left panel shows cells cultured with BrdU for 20 hours; right panel shows control cells cultured without BrdU for the same period. Gates indicate apoptotic (BrdU$^{-}$/7-AAD$^{low}$), non-dividing (BrdU$^{-}$/7-AAD$^{intermediate}$), and dividing (BrdU$^{+}$/7-AAD$^{intermediate/high}$) cell populations. (B) Cell cycle analysis of unstimulated CD4$^{+}$ T cells cultured with BrdU for 20 hours, showing the predominance of non-dividing cells. (C) Ki-67 expression in activated control CD4$^{+}$ T cells showing active proliferation. (D) Ki-67 expression in unstimulated CD4$^{+}$ T cells showing minimal proliferation.
