## Supplemental Figure 2 for "Modeling simian immunodeficiency virus (SIV) latency in primary rhesus macaque CD4^+^ T cells"

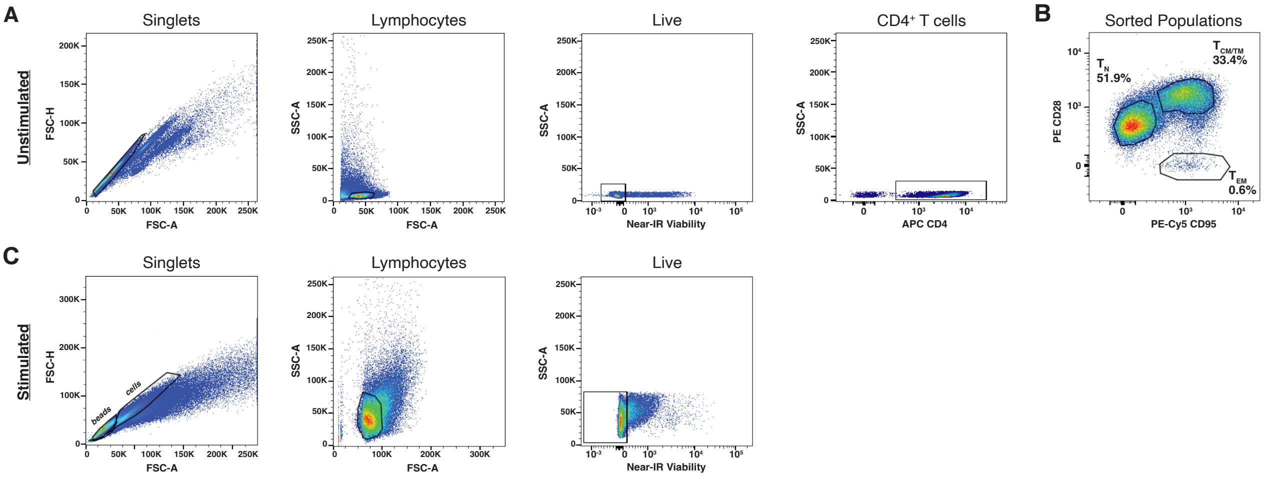


**Figure S2**. Flow cytometry gating strategies. (A) Representative flow cytometry plots showing the sequential gating strategy for identifying CD4$^{+}$ T cell subsets. Cells were gated on singlets, followed by lymphocytes, live cells, and CD4$^{+}$ cells. (B) CD4$^{+}$ T cell populations were distinguished by their expression of CD28 and CD95: naïve ($T_N$; CD28$^{+}$CD95$^{-}$), central/transitional memory ($T_{CM}$/$T_{TM}$; CD28$^{+}$CD95$^{+}$), and effector memory ($T_{EM}$; CD28$^{-}$CD95$^{+}$) populations. (C) Gating strategy for excluding T cell activation beads from analyses. Beads were identified and excluded based on their distinctive foward (FSC) and side scatter (SSC) properties, allowing for accurate analysis of cellular populations.
