## Supplemental Table 1 for "Modeling simian immunodeficiency virus (SIV) latency in primary rhesus macaque CD4^+^ T cells"

**Table S1.** Primer/probe oligonucleotide sequences for quantifying CA-vDNA by SIV *gag* ddPCR.

| **Primer/Probe** | **Sequence (5’→3’)** | **5’ Dye** | **Quencher** |
| --- | --- | --- | --- |
| SIV *gag* F primer | GTCTGCGTCATCTGGTGCATTC |  |  |
| SIV *gag* R primer | CACTAGGTGTCTCTGCACTATCTGTTTTG |  |  |
| SIV *gag* probe | CTTCCTCAGTGTGTTTCACTTTCTCTTCTG | FAM | *ZEN/IABkFQ |
| *rpp30* F primer  (single-plex) | TCAGCATGGCGGTGTTT |  |  |
| *rpp30* R primer  (single-plex) | GCTGTCTCCACAAGTC |  |  |
| *rpp30* probe  (single-plex) | TTCTGACCTGAAGGCTCTGCGC | FAM | *ZEN/IABkFQ |

** ZEN/Iowa Black FQ double quencher*
