## Supplemental Table 2 for "Modeling simian immunodeficiency virus (SIV) latency in primary rhesus macaque CD4^+^ T cells"

**Table S2**. Primer/probe oligonucleotide sequences for the SIV IPDA.

| **Primer/Probe** | **Sequence (5’→3’)** | **5’ Dye** | **Quencher** |
| --- | --- | --- | --- |
| SIV *pol* F primer | GCAGGGATAGAGCACACCTTTG |  |  |
| SIV *pol* R primer | CTATGGTTTCTACTGAATTTGCTTGTTC |  |  |
| SIV *pol* intact probe | TTTCAGGTGGTGATTCA | FAM | **MGBNFQ |
| SIV *pol* hypermutated probe | TAGGTGGTGATTTATT |  | **MGBNFQ |
| SIV *env* F primer | CCTCAATAAAGCCTTGTGTAAAATTATC |  |  |
| SIV *env* R primer | GTTGTTATTGATTTTGTCAATCCC |  |  |
| SIV *env* intact probe | TGCATTACTATGAGATGC | VIC | **MGBNFQ |
| SIV *env* hypermutated probe | TGCATTACTATAAAATGC |  | **MGBNFQ |
| SIV 2-LTR F primer | CGCCTGGTCAACTCGGTACTC |  |  |
| SIV 2-LTR R primer | GGTATGATGCCTTCTTCCTTTTCTAAG |  |  |
| SIV 2-LTR probe | CCCTGGTCTGTTAGGACCCTTTCTGCTTTG | FAM | **MGBNFQ |
| *rpp30* F primer 1 | AGGATGCTCCGGGAGTATGTA |  |  |
| *rpp30* R primer 1 | CCTGCTTGTCACCTATATAACAT |  |  |
| *rpp30* probe 1 | TCAAGCTGGGAGACGGAAGAGTCAGT | FAM | *ZEN/IABkFQ |
| *rpp30* F primer 2 | ACAGACTCACACAATTTAGG |  |  |
| *rpp30* F primer 2 | ACATTCATGCCACTGCACTC |  |  |
| *rpp30* probe 2 | ACAGGGTCTCACTTTGTTGTCCA | HEX | *ZEN/IABkFQ |

** ZEN/Iowa Black FQ double quencher*

***3’ nonfluorescent quencher/minor groove binder*
